# Characterizing the shared signals of face familiarity: long-term acquaintance, voluntary control, and concealed knowledge

**DOI:** 10.1101/2022.07.27.501568

**Authors:** Alexia Dalski, Gyula Kovács, Holger Wiese, Géza Gergely Ambrus

**Author notes:** **Corresponding author:** Géza Gergely Ambrus, Department of Psychology, Bournemouth University, Poole House, Talbot Campus, Fern Barrow, Poole, Dorset, BH12 5BB, United Kingdom.

## Abstract

Using cross-experiment multivariate classification of EEG patterns, in a recent study we found evidence for a shared familiarity signal for faces, patterns of neural activity that successfully separate trials for familiar and unfamiliar faces across participants and modes of familiarization. Here, our aim was to expand upon this research to further characterize the spatio-temporal properties of this signal. By utilizing the information content present for incidental exposure to personally familiar and unfamiliar faces, we tested how the information content in the neural signal unfolds over time under different task demands – giving truthful or deceptive responses to photographs of genuinely familiar and unfamiliar individuals. For this goal, we re-analyzed data from two previously published experiments using within-experiment leave-one-subject-out and cross-experiment classification of face familiarity. We observed that the general face familiarity signal, consistent with its previously described spatio-temporal properties, is present for long-term personally familiar faces under passive viewing, as well as for acknowledged and concealed familiarity responses. Also, central-posterior regions contain information related to deception. We propose that signals in the 200-400 ms window are modulated by top-down task-related anticipation, while the patterns in the 400-600 ms window are influenced by conscious effort to deceive. To our knowledge, this is the first report describing the representational dynamics of concealed knowledge for faces.

**Highlights:**

- Previous studies found a robust EEG effect for face-familiarity in the 200-600 ms post-stimulus range.
- This neural pattern was found to be shared across participants and modes of familiarization.
- We used incidental exposure as a template to probe information content for acknowledged and concealed familiarity
- The shared familiarity signal is modulated differentially in early (200-400 ms) and late (post-400 ms) windows
- Cross-experiment classification is a promising tool to investigate how cognitive processes unfold under different conditions

## Introduction

M/EEG studies (Dobs, Isik, Pantazis, & Kanwisher, 2019; Karimi-Rouzbahani, Ramezani, Woolgar, Rich, & Ghodrati, 2021; Wiese, Tüttenberg, et al., 2019) in the past years reported on an electrophysiological correlate of face-familiarity between 200 to 600 ms following stimulus presentation. In a recent analysis Dalski et al. (Dalski, Kovács, & Ambrus, 2022a) used data from three experiments reported in a study by Ambrus et al. (2021) to investigate whether these familiarity signals generalize across participants, stimuli, and modes of familiarization. To uncover potential shared neural signatures of face familiarity acquired through perceptual, media, and personal familiarization, a novel, cross-experiment multivariate pattern analysis (MVPA), was performed. The successful classification across experiments in the 270-630 ms time window indicated the existence of a general neural signal for face familiarity, independent of participants, familiarization methods and stimuli. Furthermore, the sustained pattern in temporal generalization analyses suggested that this signal reflects a single processing cascade.

The aim of this present paper is to expand upon these previous findings in further characterizing this general face-familiarity signal. Particularly, we are interested in how it unfolds over time under instructions to conceal familiarity with the face of a pre-experimentally highly familiar individual. This question is not only of potential importance for informing the debate over the application of brain imaging in forensic settings (Langleben & Dattilio, 2008; Langleben & Moriarty, 2013; Schauer, 2010), but can further our knowledge about the neural and cognitive processes that subserve human face perception and identification (White & Burton, 2022).

Neural processing of face familiarity has been theorized to be an automatic process. In contrast to controlled processes, automaticity entails rapid processing, even in the absence of awareness and with limited attentional resources, accompanied by a degree of independence of conscious effort and voluntary control (Yan, Young, & Andrews, 2017). Similar to not being able to “not perceive” the color of a flower as *red*, might it also be impossible not to perceive a highly familiar face as *familiar*? (Jung, Ruthruff, & Gaspelin, 2013; Ramon & Gobbini, 2018). Although much progress has been made in understanding the neural underpinnings of face familiarity and identity (Ambrus, Dotzer, Schweinberger, & Kovács, 2017; Ambrus et al., 2021; Ambrus, Kaiser, Cichy, & Kovács, 2019; Dobs et al., 2019; Nemrodov, Niemeier, Patel, & Nestor, 2018; Vida, Nestor, Plaut, & Behrmann, 2017), the nature of conscious control over these processes is far less understood. One way to test the effects of conscious control is to ask participants to suppress or modify a response to aspects of a certain stimulus (Heidlmayr, Kihlstedt, & Isel, 2020). In the case of face familiarity, this can be achieved by instructing the volunteers to actively deny knowing an otherwise familiar individual.

To date, a number of studies investigated the sequence of cognitive processing and the underlying neural basis of the active denial of familiarity with faces. An fMRI-EEG investigation by Sun et al. (Sun, Lee, & Chan, 2015) asked participants to acknowledge or deny familiarity when presented with a sequence of faces. The authors found that initial face recognition, regardless of attempts at deception, takes place at around 270 ms after stimulus onset. The two conditions diverge between 300 and 1000 ms post stimulus, indicating an effortful manipulation of information processing in this later stage. Brain areas where differential activation was found in the two conditions included the ventrolateral, dorsolateral, and dorsal medial-frontal cortices, the premotor cortex, and the inferior parietal gyrus. An fMRI imaging study by Bhatt et al. (2009) found modulation of BOLD activity in the right superior and inferior frontal gyri and bilateral precuneus for deception for familiar faces. This was further corroborated by Lee and colleagues (Lee, Leung, Lee, Raine, & Chan, 2013), who provided evidence that it is possible to detect attempts of deception by observing response pattern in the left precuneus with a high degree of accuracy.

In a recent EEG investigation, Wiese et al. (2022, Concealed Knowledge experiment) asked participants to acknowledge familiarity with a personally familiar identity, deny knowing another personally known identity, and give a truthful answer for a genuinely unfamiliar identity. This study mainly focused on two ERP components: the earlier N250 familiarity effect (Schweinberger & Neumann, 2016; Schweinberger, Pickering, Jentzsch, Burton, & Kaufmann, 2002), thought to reflect visual recognition of a known face, and the later sustained familiarity effect (SFE), hypothesized to be the marker of the integration of visual with additional identity-specific information (Wiese, Tüttenberg, et al., 2019). This study found that these components are present for both acknowledged and concealed stimuli; more specifically, measured over the TP9-TP10 and P9-P10 electrodes, they were found to be largely automatic in the ca. 200 to 400 ms time window, and modulated by attentional resources in the 400 to 600 ms interval (see also Wiese, Ingram, et al., 2019).

Despite this progress in identifying brain regions that are involved in the effortful denial of familiarity with an identity, the evolution of the representations for familiar faces under such conditions has not been investigated so far. As such, our aim here was to test if the recognition of highly personally familiar faces is indeed a single automatic processing cascade that is unaffected by active deceit. Alternatively, we wanted to probe whether the intent to deceive modulates the spatio-temporal profiles of the signals for genuinely familiar and unfamiliar faces, depending on whether familiarity is acknowledged or denied.

To answer this question, we analyzed data from two EEG-ERP experiments described in aforementioned report by Wiese et al. (2022). In the Incidental Recognition experiment, face-familiarity effects were explored using multiple images of long-term familiar and unfamiliar faces with an orthogonal target detection task. In the Concealed Knowledge experiment event-related potentials were used to determine if it is possible to detect whether a viewer is familiar with a particular face, irrespective of whether the participant is willing to acknowledge it or not.

This current study takes a data-driven approach to probe the information content present in the neural signal as it develops over time under different conditions. For this purpose, we expand upon the methods, developed by Dalski et al. (2022a, 2022b) by using multivariate cross-classification, a form of MVPA, performed across participants and experimental tasks. Instead of focusing on aggregated responses restricted in space and time, MVPA relies on the information content present in the pattern of the neural signal. In this procedure data containing neural signals is split into training and test sets, and a machine learning classifier is trained to classify test data based on the information it extracted from the training set. The classifier accuracy is therefore indicative of the presence of information in the neural signal about the categories of interest. In sum, time-resolved MVPA parses multi-dimensional, distributed patterns of neural activity to track the changes in representations relating to these categories. Here we capitalize on the methodological flexibility of MVPA to probe the evolution of active representations and cognitive processes in different contexts (Kaplan, Man, & Greening, 2015). In MVPA the train and test data need not necessarily come from the same time point, region of interest, experimental condition, participant, or experiment. Furthermore, the training and testing labels can be easily changed to test cross-classification performance across different tasks and domains. This remarkable flexibility allows us to establish a template, a sort of “ground truth”, where familiar face perception unfolds unimpeded. This data can then be utilized to investigate the changes in the evolution of information processing under differential task demands and contexts.

While previous studies used univariate analyses (e.g., Bhatt et al., 2009; Lee et al., 2013; Sun et al., 2015) and within-participant classification (Wiese et al., 2022) to investigate the effects of conscious control over the neural underpinnings of face recognition, this study aims at probing the development of the information content in the neural signal under specific task conditions, in relation to each other.

In order to achieve this objective, we needed to make certain that a number of prerequisites are met. The most important requirement to test cross-participant and cross-task classification is the existence of a shared face-familiarity signal for long-term acquaintance. While Dalski et al. (2022a) found high cross-participant and cross-experiment decodability for the personally familiarized faces, the familiarization phase was intentionally kept uniform for all participants. This standardized experimental familiarization regime was useful to assess differences between pre- and post-familiarization neural signals, but it left open the question of their development in more organic, long-term settings. On the one hand, several studies argue for the importance of affective, semantic, episodic information in the development of face-familiarity signals underlying the cognitive/neural processing of long-term personal familiarity (Campbell & Tanaka, 2021; Popova & Wiese, 2022; Wiese et al., 2022, for a review, see Kovács, 2020). Thus, as a result of the accumulation of diverse experiences that are unique to each dyad, theoretically, it is possible that every person has a distinctive neural code for representing long-term personally familiar identities. On the other hand, Wiese et al. (2021b) found that familiarity-related ERP components for familiar faces are quantitatively, but not qualitatively modulated by the depth of familiarity. Furthermore, a recent fMRI hyper-alignment study by Visconti di Oleggio Castello et al. (2021) demonstrated that individually distinctive information associated with familiar faces is embedded in a neural code that is shared across brains. Finally, Dalski et al. (Dalski, Kovács, & Ambrus, 2022b) had shown that shared neural signatures of face familiarity can arise without any additional semantic, affective, or social information, arguing for a broad generalizability of the signal. Nevertheless, the existence of the general neural signature for long-term, pre-experimental familiarity for faces has not yet been established. One aim of this study is therefore to formally test this assumption.

Another important factor relates to the robustness of the signal under different task instructions. Again, Dalski et al. (2022b) demonstrated that the shared face-familiarity signal can be observed under markedly different task conditions (successful cross-classification between passive exposure to single face stimuli versus a face-matching task with multiple faces presented simultaneously). Nevertheless, while Dalski et al. required the participants to be as accurate as possible with their responses, the Concealed Knowledge study analyzed in this report asked participants to actively suppress the truthful answers regarding the target stimuli. Thus, establishing the cross-participant and cross-dataset decodability of familiarity for familiar faces with active denial of their knowledge is not only of theoretical importance, but a precondition for further investigation into the characteristics of the general familiarity signal and its evolution.

In summary, this present study aims to test the following hypotheses: 1) Can the general signal of face familiarity reported in previous studies for short-term, experimentally familiarized faces be also observed for long term, participant-unique personally familiar faces? 2) Is this signal indicative of an automatic process, independent from voluntary control and attempts of deceit? 3) Is there a difference in the spatio-temporal profiles indicative of deception when exposed to personally familiar faces?

## Methods

### Datasets

The time-resolved within- and cross-experiment generalization analyses were carried out using data from the Incidental Recognition (n=22) and Concealed Knowledge (n=19) experiments reported in Wiese et al. (2022). Participants gave written, informed consent, and the study was approved by the ethics committee of Durham University’s Psychology Department.

Stimuli in both experiments consisted of photos of unknown and highly personally familiar faces (e.g. close friends, relatives) that were cropped to 190 × 285 pixels, converted to grayscale, and luminance-adjusted using the SHINE toolbox (Willenbockel et al., 2010). The images were presented for 1000 ms on a gray background. The duration of the inter-stimulus interval (with fixation cross) varied randomly between 1500 and 2500 ms.

The original aim of the Incidental Recognition experiment was to test the difference in ERPs between high- and low variability stimuli (multiple vs. single photographs of known and unknown identities), thus the stimuli consisted of photographs of four identities, two pairs of a known and an unknown person, one pair presented with multiple images (trial-unique, high variability), the other with a single image, repeated throughout the experiment (no variability). In the interest of comparability, we used the high variability familiar and high variability unfamiliar trials of the incidental exposure experiment. These conditions will be referred to as (incidental) familiar and unfamiliar in the remainder of this paper.

The Concealed Knowledge experiment tested the effect of attentional load and voluntary control on familiarity-related ERP components. In this experiment trial-unique images of one unfamiliar and two familiar persons were presented. The participants were instructed to truthfully indicate familiarity with the unfamiliar and one of the familiar faces (unfamiliar and acknowledged familiar) and deny familiarity with (i.e., respond ‘unfamiliar’ to) the other familiar identity (concealed familiar).

While the Incidental Recognition experiment required no identity-related response from the participants (no response button press during trials with a face being presented; key press with right hand index finger to trials with butterflies), the Concealed Knowledge experiment directly required a response corresponding to the face-image being presented (left and right index fingers, counterbalanced across participants).

### Within-experiment classification of familiarity

As an initial step to establish if the prerequisites are met for our main objective, we conducted a within-experiment (leave-one-subject-out) decoding analysis. This was performed in order to test two of our hypotheses: 1) to establish whether neural patterns for participant-unique, highly personally familiar faces can be cross-classified; and 2) if deception can be detected – i.e., if information in the neural patterns in ERPs for acknowledged and concealed trials can be used to successfully separate these signals across participants. Time resolved decoding (Grootswagers, Wardle, & Carlson, 2017), using Linear Discriminant Analysis (LDA) classifiers, was performed on all electrodes and pre-defined regions of interest (left and right anterior, central, and posterior regions) (**Figure 1**, left panel).

**Figure 1.**
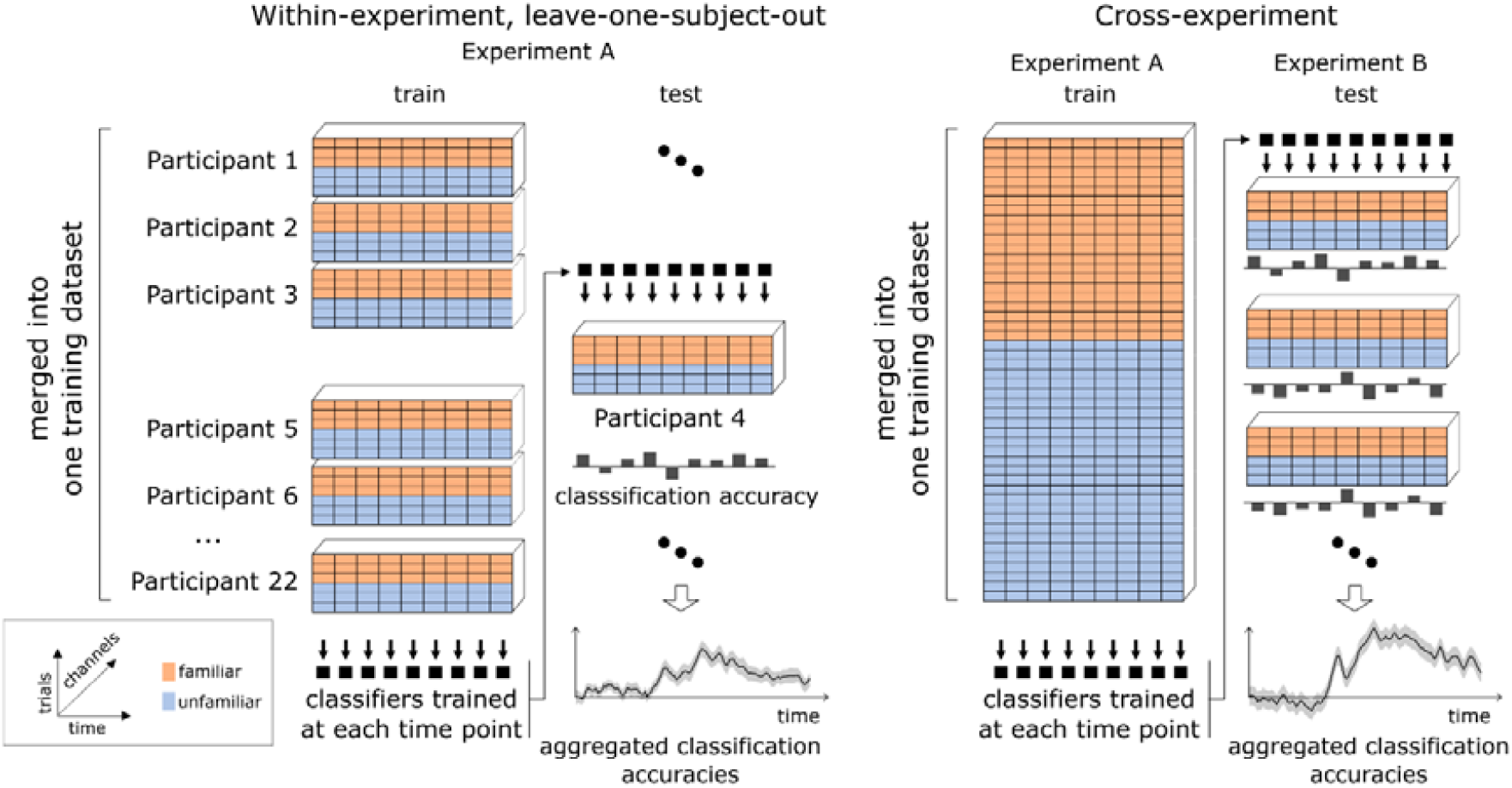
Analysis pipelines. In within-experiment leave-one-subject-out cross-validation (left) the data of each participant is iteratively left out for testing, while data from the remaining participants is collated into the test set. Classification accuracies for each participant are then aggregated. In cross-experiment classification (right) classifiers are trained only once on collated data from all participants in one experiment; these classifiers are then used to predict class membership for data from another experiment, for each participant separately. These individual decoding accuracies are then aggregated on a sample level.

### Cross-experiment classification of familiarity

To further test whether the neural signatures for highly familiar identities generalize across participants and conditions depending on attempts at deception, we also performed cross-experiment classification analyses based on these two experiments in both directions (**Figure 1**, right panel). For this purpose, time-resolved decoding was performed over all electrodes and pre-defined regions of interest. The analysis pipeline was identical to the one described in Dalski et al. (Dalski et al., 2022a). The procedures performed are summarized in **Table 1**.

**Table 1.** Summary of the cross-experimental decoding analyses. For each analysis, both in the Forward and Reverse directions, labels for the Incidental Exposure experiment remained the same (Familiar and Unfamiliar). In the case of the Concealed Knowledge experiment, labels for classifier training and testing were replaced according to the research question.

|  | Training | Condition | New label | Test | Condition | New label |
| --- | --- | --- | --- | --- | --- | --- |
| Forward<br>Acknowledged | Incidental<br>Knowledge | Familiar | *Familiar* | Concealed<br>Knowledge | Acknowledged Familiar | *Familiar* |
|  |  | Unfamiliar | *Unfamiliar* |  | Unfamiliar | *Unfamiliar* |
| Forward<br>Concealed | Incidental<br>Recognition | Familiar | *Familiar* | Concealed<br>Knowledge | Concealed Familiar | *Familiar* |
|  |  | Unfamiliar | *Unfamiliar* |  | Unfamiliar | *Unfamiliar* |
| Forward Lie | Incidental<br>Recognition | Familiar | *Familiar* | Concealed<br>Knowledge | Acknowledged Familiar | *Familiar* |
|  |  | Unfamiliar | *Unfamiliar* |  | Concealed Familiar | *Unfamiliar* |
| Reverse<br>Acknowledged | Concealed<br>Knowledge | Acknowledged Familiar | *Familiar* | Incidental<br>Recognition | Familiar | *Familiar* |
|  |  | Unfamiliar | *Unfamiliar* |  | Unfamiliar | *Unfamiliar* |
| Reverse<br>Concealed | Concealed<br>Knowledge | Concealed Familiar | *Familiar* | Incidental<br>Recognition | Familiar | *Familiar* |
|  |  | Unfamiliar | *Unfamiliar* |  | Unfamiliar | *Unfamiliar* |
| Reverse Lie | Concealed<br>Knowledge | Acknowledged Familiar | *Familiar* | Incidental<br>Recognition | Familiar | *Familiar* |
|  |  | Concealed Familiar | *Unfamiliar* |  | Unfamiliar | *Unfamiliar* |

In the *Forward* direction, classifiers were trained on ERPs from the Incidental Recognition experiment and tested on those of the Concealed Knowledge study. This means, ERP data was collated across participants in the Incidental Recognition, and classifiers were trained at each time-point to differentiate between Familiar and Unfamiliar trials. These classifiers then were used to predict in the Concealed Knowledge (a) *Acknowledged* familiarity: the accuracy in classifying Acknowledged Familiar and Unfamiliar trials; (b) *Concealed* familiarity: the accuracy in differentiating between Concealed Familiar and Unfamiliar trials.

In the *Reverse* direction, classifiers were trained on ERPs from pairs of conditions in the Concealed Knowledge experiment (collated across participants) and tested on those of the Incidental Recognition study. Briefly, classifier accuracies at each time-points were tested when training was performed on (a) *Acknowledged* Familiarity and Unfamiliar trials; (b) *Concealed* Familiarity and Unfamiliar trials.

The Acknowledged condition was devised to test if information that is useful to classify familiar and unfamiliar face stimuli ERPs in Incidental Recognition can separate acknowledged familiar and unfamiliar trials, and vice versa. The Concealed condition tested if true familiarity can be cross-classified, irrespective of attempts at deception in the Concealed Knowledge experiment. In other words, these conditions test our hypothesis whether familiarity signals generalize across participants and experimental designs.

### The modulation of familiarity signals by deliberate deception

We devised one further test of the modulation of the general familiarity signal by deception. The *Lie* condition was designed to assess the extent by which the intent of deception can influence cross-classification accuracies towards the deceptive answer. This required the relabeling of the concealed familiarity condition of the Concealed Knowledge experiment for the purposes of classifier training and testing, i.e., accepting the deceptive answer, differentiating between acknowledged familiar and concealed familiar trials by treating the latter as if these were unfamiliar.

### Statistical testing

In order to increase the signal-to-noise ratio, all analyses were conducted on data downsampled to 100 Hz, and a moving average of 30 ms (3 consecutive time-points) was applied to all within-experiment and cross-experiment decoding accuracy data at the participant level (Ambrus et al., 2019; Dalski et al., 2022a; Kaiser, Oosterhof, & Peelen, 2016). For the time-resolved ROI-based analyses, decoding accuracies were entered into two-tailed, one-sample cluster permutation tests (10,000 iterations) against chance (50%). We also tested cross-experiment decoding accuracies across 6 train/test pairings with two-tailed cluster permutation tests with 10,000 iterations.

The classification and the statistical analyses were conducted using python, MNE-Python (Gramfort et al., 2013), scikit-learn (Pedregosa et al., 2011) and SciPy (Virtanen et al., 2020).

## Results

### Within experiment cross-classification

In the within-experiment, leave-one-subject-out classification results we see robust familiarity effects for incidental, acknowledged, and concealed familiarity, when tested against ERPs for unfamiliar stimuli, with sustained significant clusters starting around 200 ms in all regions of interest (see **Supplementary Table 1**. for detailed statistics). This supports our hypotheses that familiarity with highly personally familiar faces can be cross-classified across participants, even when the identities are unique to each volunteer.

On the other hand, cross-participant decodability between concealed and acknowledged familiarity was restricted to a brief period between ca. 200-330 ms over the right central and bilateral posterior ROIs. This means that this early time window that contains information separating actual familiarity and unfamiliarity with faces, also holds information that separates concealed and acknowledged familiarity.

### Cross-experiment classification

#### Genuine familiarity independent of deception

Results of the time-resolved cross-experiment classification are depicted in **Figure 3**. All-electrodes and region-of-interest-based decoding analyses, in both Forward, and Reverse directions, show strong, significant cross-experiment decodability of actual face familiarity, starting around 200 ms, and lasting up to 800 ms post-stimulus onset, and beyond (see **Supplementary Tables 2**. for detailed statistics). This confirms two of our hypotheses: 1) incidental familiarity and acknowledged familiarity can be cross-classified regardless of differences in task in the two experiments; and 2) the limited conscious control over the shared familiarity signal is indicated by the successful cross-classifiability of incidental and concealed familiarity, against unfamiliar trials.

#### The effects of deception

The Lie condition was designed to test if participants’ intent to deceive indeed leads to neural patterns that resemble genuine unfamiliarity; in other words, whether classifiers could be “deceived” into accepting the deception and cross-classify patterns for genuinely familiar and unfamiliar faces correctly.

While we found no significant clusters for the Lie condition neither in the Forward, nor the Reverse decoding direction over all electrodes, interestingly, strong, significant clusters emerged in all ROIs, except for the right posterior region, between ca. 400 and 600 ms in the reverse Lie condition. This was most prominent over the bilateral central regions of interest. Furthermore, in both Forward and Reverse directions, a negative cluster between 200 and 400 ms emerged in the right posterior ROI (**Figure 4**., see **Supplementary Table 2** for detailed statistics).

#### The evolution of representations for different stimulus categories

Our main analysis follows up this interesting result by probing the information content present for the three types of stimuli in the Concealed Knowledge experiment. For that purpose, we used the Incidental Recognition data as the “ground truth”, i.e., the “canonical” time course of the evolution of the representations for familiar and unfamiliar faces. First, we separately characterized the classification accuracy profiles of familiar and unfamiliar face stimuli in the within-experiment, leave-one-subject-out analysis (**Figure 5**., **left panel**). This demonstrated that for passive viewing of faces, information that identifies familiar and unfamiliar trials is present and sustained from ca. 200 ms after stimulus onset. To examine how this information is then utilized to cross-classify true unfamiliarity, acknowledged and concealed familiarity, we examined cross-experiment classification accuracies in the Forward direction for these stimuli separately as well (**Figure 5**., **right panel**).

While the cross-classification accuracy for the true unfamiliar signals, starting at ca. 200 ms, remained constant and robust throughout much of the epoch, we saw a dissociation between concealed and acknowledged familiarity in the early (200-400 ms) and late (post-400 ms) time windows. Overall, the difference in cross-classification accuracies between these two conditions were not shown to be significant in any of the ROIs. However, over all electrodes and bilateral central and posterior sites, clusters flagged as significant for concealed trials fell into the early window, while significant clusters for acknowledged trials were present in the later period. Furthermore, in these regions of interest and time windows, significant clusters emerged when comparing both acknowledged and concealed familiar to true unfamiliar trials (see **Supplementary Table 3**. for detailed statistics).

## Discussion

The objective of this study was to investigate how the neural representations of familiarity unfold over time under different task demands, specifically for acknowledging and denying familiarity with personally familiar individuals. For this purpose, we related signals for unfamiliar, acknowledged, and concealed familiar faces to signals for incidental recognition of familiar and unfamiliar faces by means of cross-participant and cross-experiment classification. Our main findings were the following: 1) Highly personally familiar, participant-unique faces elicit a shared, general familiarity signal that differentiates between familiar and unfamiliar identities across participants and tasks. This signal is preserved even when participants are instructed to conceal knowledge of a familiar identity. 2) This signal can be successfully utilized as a template across experiments to interrogate the evolution of face familiarity processing under different task demands. 3) A clear dissociation between acknowledged and concealed familiarity emerges between early (ca. 200-400 ms) and late (post-400 ms) processing stages.

### Cross-Experiment Face Familiarity Decoding Replicates for Long-term Familiarity

In Dalski, Kovács, & Ambrus (Dalski et al., 2022a) we reported on the generalizability of face-familiarity signals across experiments. The data for that study came from three experiments reported in Ambrus, Eick, Kaiser, & Kovács (Ambrus et al., 2021), where experimental familiarization was achieved either perceptually, via media exposure, or by personal interaction. We observed significant cross-experiment familiarity decoding involving all three experiments, predominantly over posterior and central regions of the right hemisphere in the 270–630 ms time window. This study replicates this finding on an independent dataset. In our present study, acknowledged/unfamiliar trial ERPs were decodable in both Forward and Reverse decoding directions (i.e., training on the Incidental Recognition and testing on the Concealed Knowledge experiment, and vice versa) starting from around 200 ms, and extending to 800 – 1000 ms post-stimulus onset (**Figure 3**., **blue lines**). The spatio-temporal profile of the decoding accuracy time-course is comparable to that reported in Dalski, Kovács, & Ambrus (Dalski et al., 2022a), providing further support for the existence of a robust, general face-familiarity signal. Here, this signal is demonstrated to be present for pre-experimental, long-term personal familiarity, and across experiments that used individualized, trial-unique stimuli, and required differential engagement with the images. Establishing cross-participant and cross-task decodability for participant-unique, personally highly familiar faces allowed us to further test the characteristics of this general familiarity signal under different task demands.

### Concealed and Acknowledged Knowledge Elicit Comparable General Familiarity Signals

When tested against ERPs elicited by true unfamiliar faces, acknowledged, and concealed familiar faces elicited comparable general familiarity signals across all regions of interest (**Figure 3**., **green lines**). Formal statistical testing found no substantial differences in the cross-decoding accuracy time-courses in any of the regions of interest on the sample level, neither in the Forward nor the in the Reverse direction, indicating that the general face-familiarity signal survives attempts of deception. The implications of this finding for the detection of concealed knowledge are discussed in the following sections.

### Differentiating Concealed and Acknowledged Familiarity

The within-experiment, concealed vs. acknowledged analysis (**Figure 2, red line**) was conducted to test whether classifiers can extract information indicative of the intention to deceive, when participants view highly personally familiar faces. The results of this procedure revealed an early (ca. 200-330 ms) central-posterior effect, coinciding with the approximate onset of the effects observed for incidental, acknowledged, and concealed familiarity. This indicates that information across participants’ neural patterns exists that differentiates the processing of two otherwise familiar faces on the basis of whether familiarity with the given faces can be freely acknowledged or needs to be denied. That this effect is solely due to the intention to deceive and not related to factors linked to image or identity, is supported by the fact that the familiar face stimuli were unique to each participant, and the to-be-acknowledged and to-be-concealed identities were chosen randomly.

**Figure 2.**
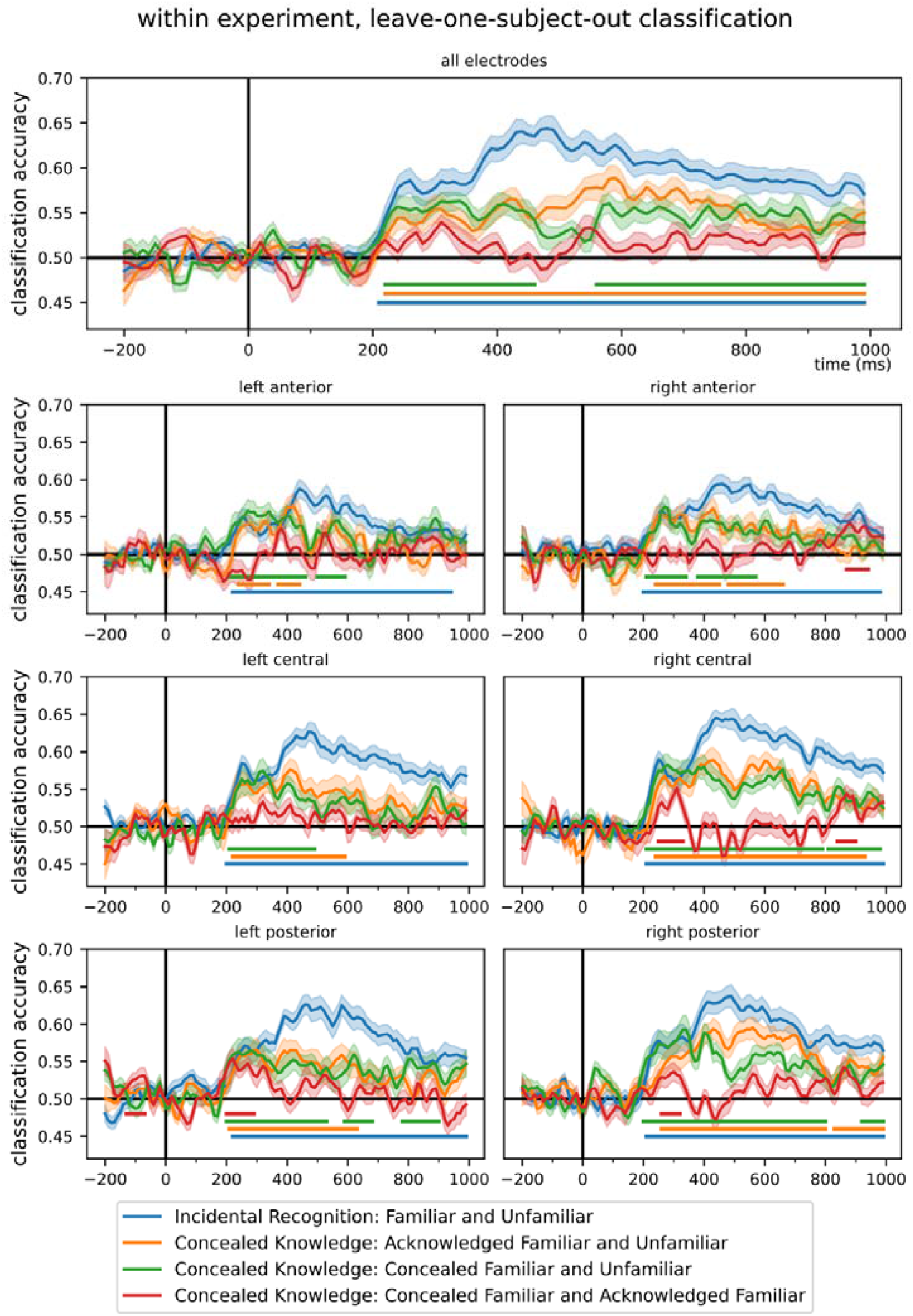
Within-experiment, time-resolved, leave-one-subject-out classification. The classifiers trained on same-experiment data were able to decode genuine face-familiarity with a high degree of accuracy starting around 200 ms after stimulus onset. The classifiers trained to differentiate between signals for concealed familiar and acknowledged familiar faces were only able to successfully do so in the left central and bilateral posterior ROIs (Ambrus et al., 2019) between ca. 200-400 ms. Shaded ranges denote ±SEM. For detailed statistics see **Supplementary Table 1**.

While for incidental, acknowledged, and concealed familiarity we observed a sustained pattern of cross-participant within-experiment classifiably lasting several hundred milliseconds after stimulus onset (**Figure 2**., **blue, green, and orange lines**), the concealed vs. acknowledged effect was restricted to this initial time window, and to the regions of interest where otherwise face-familiarity effects are found to be the most prominent.

When contrasting concealed and acknowledged familiarity directly by re-labeling the concealed trials as “unfamiliar” (consistent with the response the participants were required to give, Lie condition) two observations became apparent. 1) Over all electrodes, within-experiment and cross-experiment classification was not able to differentiate between the two conditions. 2) Over central and posterior electrode clusters a dissociation between earlier (ca. 200-400 ms) and later (ca. 400-700 ms) phases, was observed (**Figure 4**.).

Corresponding to the early (220-360 ms) within-experiment effect described above, we found significant right posterior decodability in the cross-experiment Lie condition, interestingly, in the *negative* direction: concealed trials were *more likely* to be classified as familiar than acknowledged trials). While at this stage we have no definitive explanation for this, one possibility is that as a preparation to give the expected deceptive response to faces in the concealed condition, representations are already active in anticipation of seeing the concealed identity. Thus, when to-be-concealed faces do get presented, the visual input may reinforce the already active representation, leading to an enhanced early familiarity signal. Indeed, task-related modulation of informative neural response patterns has been reported (Bonte, Hausfeld, Scharke, Valente, & Formisano, 2014; Yip, Cheung, Ngan, Wong, & Wong, 2022), and it has been previously shown that task context selectively enhances the processing of task-relevant stimuli (Grootswagers, Robinson, Shatek, & Carlson, 2021).

Starting from 380 ms, most consistently over left and right central clusters, classifiers trained on the concealed and acknowledged trials in the Concealed Knowledge experiment were able to differentiate between familiar and unfamiliar trials in the Incidental experiment (Reverse *Lie*). This may indicate that either the intention to deceive, or the increased effort to suppress a truthful answer, shifted the patterns to the unfamiliar direction sufficiently enough to allow for successful cross-classification (**Figure 5**., **green line**).

Based on these findings, it is therefore conceivable that both early and late signals in these regions of interest are susceptible to task-related modulations. The early (200-400 ms) window might be influenced by top-down effects of task relevance, and later (post-400 ms) signals by bottom-up conscious control. More research is needed to replicate and further investigate these findings, as if this pattern is found to be robust in the future, it may be a potential biomarker indicative of deception.

### Potential to Reveal Concealed Knowledge

Compared to more widely used within-participant cross-validation methods, cross-participant and cross-experiment decoding benefits from the availability of a larger/more diverse sets of training data. The training set in such a design is a combination of data from several participants, the exemplars are more varied, leading to the reduction of the effects arising from idiosyncratic participant-level and stimulus properties. In our present analysis it allowed us to investigate neural signals for genuinely familiar faces under instructions to conceal familiarity with a given identity. We found an early, cross-participant effect indicative of deception, which we theorize to be the result of anticipation to give a deceptive answer to the face of a given familiar identity. Decoding familiarity/unfamiliarity from all sensors, cross-experiment classification found no substantial differences in the cross-classification profiles for concealed and acknowledged familiarity (**Figure 3**), and the effects of concealment were also restricted to specific ROIs when the classifiers were fed deceptive labels (**Figure 4**). Of particular note, the time-course of cross-classification for true unfamiliar trials were largely unaffected (**Figure 5**., **left, grey lines**). This observation may have practical implications in establishing “innocence”, i.e., signals for true unfamiliar faces are likely to be detected. Future studies are needed to test this hypothesis, and under what conditions it might hold true – for example giving a deceptive answer about knowing a genuinely unfamiliar identity.

**Figure 3.**
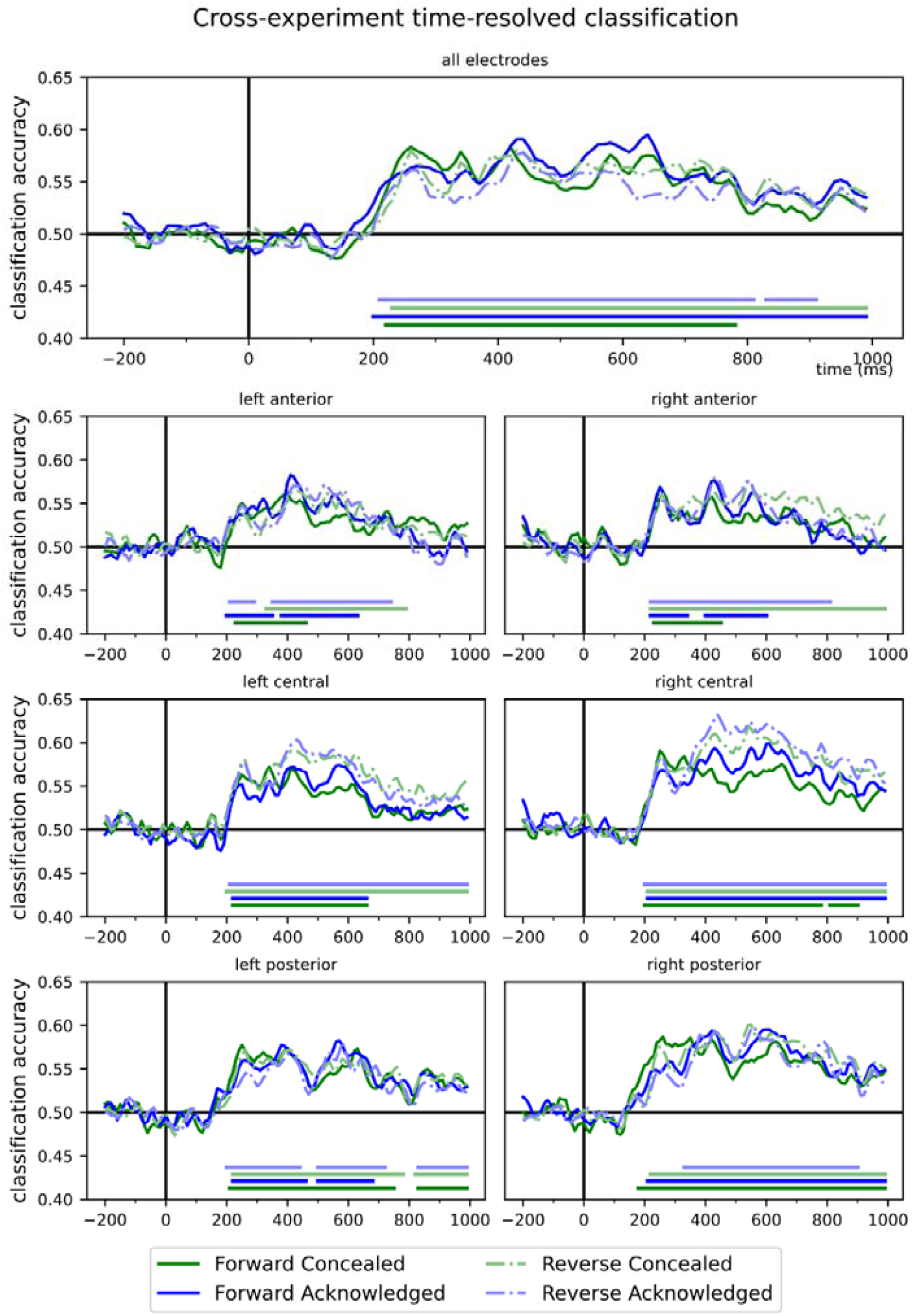
Time-resolved cross-experiment classification. Forward: Training on Incidental Recognition, testing on Concealed Knowledge. Reverse: training on Concealed Knowledge, testing on Incidental Recognition. For *acknowledged*/true unfamiliar and *concealed*/true unfamiliar, significant cross-experimental decodability was observed ca. 200 ms following stimulus onset in both Incidental to Concealed (Forward) and Concealed to Incidental (Reverse) directions (blue and green lines). (Two-tailed cluster permutation tests.) For detailed statistics see **Supplementary Table 2, a-d**.

**Figure 4.**
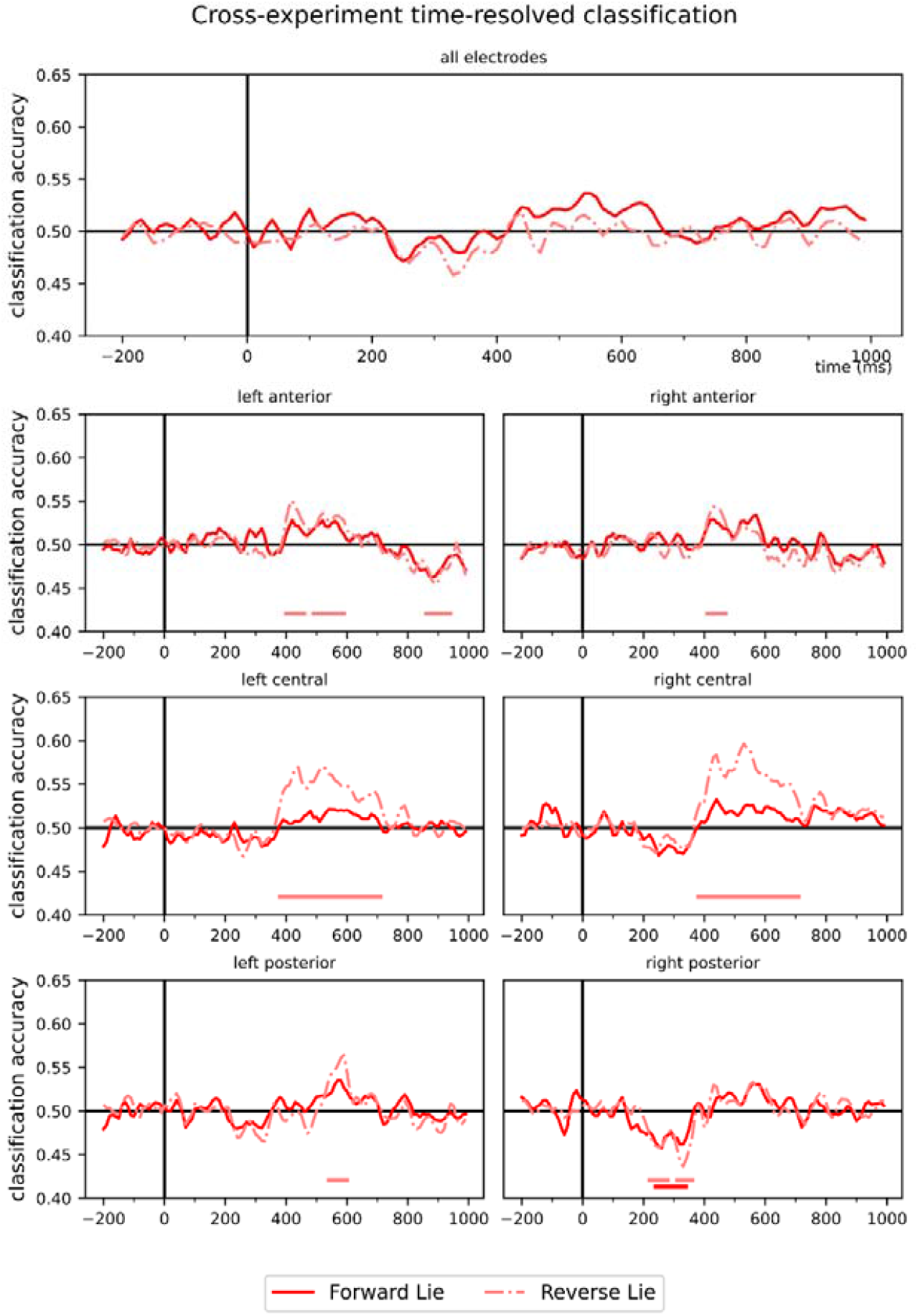
Time-resolved cross-experiment classification for acknowledged and concealed trials, with concealed trials relabeled as unfamiliar (Lie condition). In both Forward and Reverse directions significant negative clusters over the right posterior ROI between 220 - 360 ms emerged, indicating that the classifiers treated the concealed trials as *more familiar*. In the Reverse direction only, most prominently over both left and right central sites, positive clusters were seen between 380 and 710 ms, indicating that in this time window and ROIs, the classifiers treated the concealed trials as *more unfamiliar*. No significant clusters were found over all electrodes in either cross-decoding direction. Forward: Trained on Incidental (familiar vs. unfamiliar) tested on Concealed (concealed as unfamiliar vs. familiar), Reverse: Trained on Concealed (concealed as unfamiliar vs. familiar), tested on Incidental (familiar vs. unfamiliar). Statistics were calculated using two-tailed cluster permutation tests (p<0.05). For detailed statistics see **Supplementary Table 2, e-f**.

**Figure 5.**
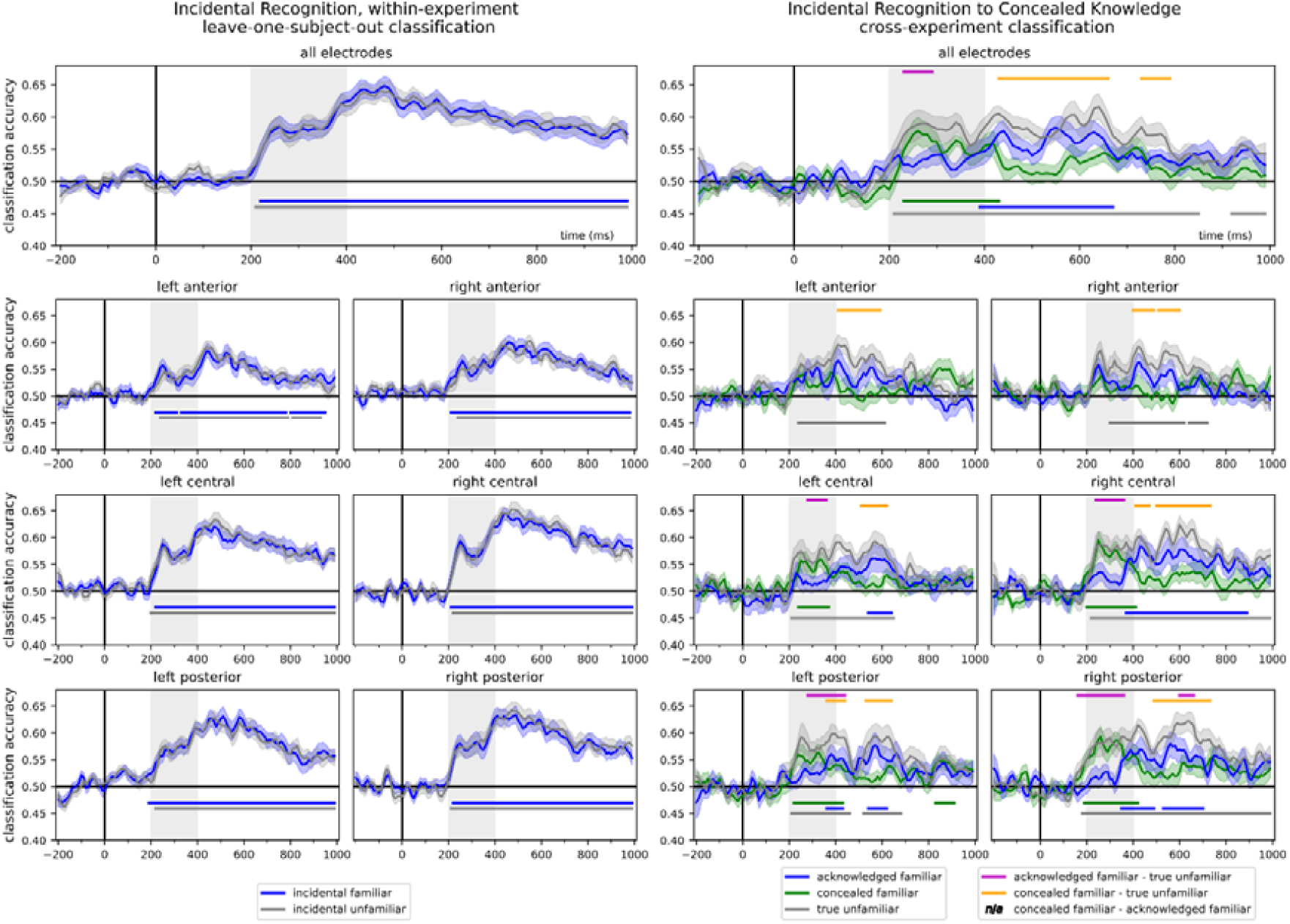
Tracking the evolution of information content in the EEG signal. Time-resolved classification accuracies shown separately for the different stimulus categories in the (left) Incidental Recognition experiment (within-experiment, leave-one-subject-out pipeline) and the (right) cross-experiment decoding in the Forward (Incidental to Concealed Knowledge) direction. Results of the LOSO decoding in the Incidental Recognition experiment show that decoding accuracies do not differ between ERPs for familiar and unfamiliar faces, and information that can aid the classification of familiar and unfamiliar conditions is present in the data from around 200 ms following stimulus onset (**Supplementary Table 3**., **a-b**). However, when using this information to decode concealed and acknowledged familiarity in the Concealed Knowledge experiment, a clear separation at ca. 400 ms was observed. While the cross-experiment decoding of the true unfamiliar condition remained largely unaffected, classifier accuracies for the concealed familiar condition peaked earlier (around 200 to 400 ms, marked for clarity), significant clusters for the acknowledged familiar condition started to appear only after around 400 ms post stimulus onset **(Supplementary Table 3**., **c-e**)., reflecting also in significant differences when compared to the unfamiliar condition in these time windows (**Supplementary Table 3**., **f-g**). No difference between acknowledged and concealed familiarity decoding accuracies was observed. Shaded ranges denote ±SEM. Statistics were calculated using two-tailed cluster permutation tests.

### Summary

In summary, we have demonstrated that cross-experiment classification can be a useful tool to investigate how cognitive and neural processes unfold over time under different task demands. Here, the information content in the EEG patterns for incidental exposure to familiar and unfamiliar faces was used to characterize the representational dynamics of acknowledged, concealed, and unfamiliar faces. Using this method, we have shown that general, shared face familiarity signals are modulated differentially in early (ca. 200-400 ms) and late (post-400 ms) time windows, likely reflecting top-down and bottom-up processing related to task demands. We hope to see the exploration of the usefulness of this method in other domains both in basic and in applied research.

## List of Supplementary Information

**Supplementary Table 1**. Statistics - Within-experiment, leave-one-subject-out, time-resolved cross-classification

**Supplementary Table 2**. Statistics – Cross-experiment classification

**Supplementary Table 3**. Statistics – Classification of stimulus types separately; Incidental Recognition, within-experiment, Concealed Knowledge, cross-experiment, Forward direction.

