## Supplementary Table 1 for "Characterizing the shared signals of face familiarity: long-term acquaintance, voluntary control, and concealed knowledge": SupplementaryTable_1.html

  

a) **Incidental Recognition: incidental familiar vs. incidental unfamiliar**

|  |  |  |  |  |  |  |  |  |
| --- | --- | --- | --- | --- | --- | --- | --- | --- |
|  | time window | peak latency | cluster *p* | peak Cohen's *d* |  | | | |
| **all electrodes** | 210 - 990 ms | 480 ms | 0.0001 | 2.1658 |  | | | |
|  | **left hemisphere** | | | | **right hemisphere** | | | |
|  | time window | peak latency | cluster *p* | peak Cohen's *d* | time window | peak latency | cluster *p* | peak Cohen's *d* |
| **anterior** | 220 - 940 ms | 440 ms | 0.0001 | 1.575 | 200 - 980 ms | 460 ms | 0.0001 | 1.6355 |
| **central** | 200 - 990 ms | 470 ms | 0.0001 | 2.242 | 210 - 990 ms | 440 ms | 0.0001 | 2.9577 |
| **posterior** | 220 - 990 ms | 460 ms | 0.0001 | 2.016 | 210 - 990 ms | 490 ms | 0.0001 | 1.9183 |

  

b) **Concealed Knowledge: acknowledged familiar vs. true unfamiliar**

|  | time window | peak latency | cluster *p* | peak Cohen's *d* |  | | | |  |
| **all electrodes** | 220 - 990 ms | 590 ms | 0.0001 | 1.4765 |  | | | |  |
|  | | | | | | | | |  |
|  | **left hemisphere** | | | | **right hemisphere** | | | |  |
|  | time window | peak latency | cluster *p* | peak Cohen's *d* | time window | peak latency | cluster *p* | peak Cohen's *d* |  |
| **anterior** | 240 - 340 ms | 330 ms | 0.0128 | 0.7012 | 240 - 450 ms | 430 ms | 0.0009 | 0.9866 |  |
 370 - 440 ms | 420 ms | 0.024 | 0.9436 | 480 - 660 ms | 560 ms | 0.0061 | 0.6299 ||  | **central** | 220 - 590 ms | 410 ms | 0.0004 | 1.2292 | 240 - 930 ms | 430 ms | 0.0001 | 1.2366 |
| **posterior** | 210 - 630 ms | 300 ms | 0.0004 | 0.981 | 260 - 800 ms | 560 ms | 0.0001 | 1.4924 |  |
  | | | | 830 - 990 ms | 990 ms | 0.0193 | 0.8362 |

  

c) **Concealed Knowledge: concealed familiar vs. true unfamiliar**

|  | time window | peak latency | cluster *p* | peak Cohen's *d* |  | | | |  |
| **all electrodes** | 220 - 460 ms | 340 ms | 0.0011 | 1.622 |  | | | |  |
 560 - 990 ms | 580 ms | 0.0001 | 1.2712 |  | | | ||  |  | | | | | | | | |
|  | **left hemisphere** | | | | **right hemisphere** | | | |  |
|  | time window | peak latency | cluster *p* | peak Cohen's *d* | time window | peak latency | cluster *p* | peak Cohen's *d* |  |
| **anterior** | 210 - 460 ms | 270 ms | 0.0003 | 0.8984 | 210 - 340 ms | 270 ms | 0.0053 | 0.9645 |  |
 500 - 590 ms | 570 ms | 0.0348 | 0.8062 | 380 - 570 ms | 420 ms | 0.0009 | 1.0703 ||  | **central** | 210 - 490 ms | 320 ms | 0.0016 | 1.0387 | 210 - 790 ms | 280 ms | 0.0001 | 1.294 |
  | | | | 810 - 980 ms | 840 ms | 0.011 | 1.0542 || **posterior** |  | 200 - 530 ms | 280 ms | 0.0012 | 0.8297 | 200 - 800 ms | 330 ms | 0.0001 | 1.1635 |
 590 - 680 ms | 620 ms | 0.0414 | 0.9476 | 920 - 990 ms | 990 ms | 0.027 | 0.732 | 780 - 900 ms | 810 ms | 0.0172 | 1.0992 |  | | | |

  

d) **Concealed Knowledge: concealed familiar vs. acknowledged familiar**

|  | time window | peak latency | cluster *p* | peak Cohen's *d* |  | | | |  |
| **all electrodes** |  | | | |  | | | |  |
|  | | | | | | | | |  |
|  | **left hemisphere** | | | | **right hemisphere** | | | |  |
|  | time window | peak latency | cluster *p* | peak Cohen's *d* | time window | peak latency | cluster *p* | peak Cohen's *d* |  |
| **anterior** |  | | | | 870 - 940 ms | 880 ms | 0.0389 | 0.6756 |  |
| **central** |  | | | | 250 - 330 ms | 310 ms | 0.0063 | 1.2417 |  |
  | | | | 840 - 900 ms | 870 ms | 0.0161 | 0.8469 ||  | **posterior** | -130 - -70 ms | -80 ms | 0.0466 | 0.6388 | 260 - 320 ms | 300 ms | 0.0441 | 0.6904 |
 200 - 290 ms | 220 ms | 0.0021 | 0.8798 |  | | | |

  
