## Supplementary Table 2 for "Characterizing the shared signals of face familiarity: long-term acquaintance, voluntary control, and concealed knowledge": SupplementaryTable_2.html

  

a) **Forward Acknowledged**  
trained on Incidental Recognition: incidental familiar vs. incidental unfamiliar  
tested on Concealed Knowledge: acknowledged familiar vs. true unfamiliar

|  | time window | peak latency | cluster *p* | peak Cohen's *d* |  | | | |  |
| **all electrodes** | 200 - 990 ms | 640 ms | 0.0001 | 1.4147 |  | | | |  |
|  | | | | | | | | |  |
|  | **left hemisphere** | | | | **right hemisphere** | | | |  |
|  | time window | peak latency | cluster *p* | peak Cohen's *d* | time window | peak latency | cluster *p* | peak Cohen's *d* |  |
| **anterior** | 200 - 350 ms | 320 ms | 0.0135 | 1.121 | 220 - 340 ms | 250 ms | 0.0174 | 1.4207 |  |
 380 - 630 ms | 410 ms | 0.0022 | 1.0045 | 400 - 600 ms | 430 ms | 0.0023 | 1.3652 ||  | **central** | 220 - 660 ms | 570 ms | 0.0005 | 0.8621 | 210 - 990 ms | 610 ms | 0.0001 | 1.3339 |
| **posterior** | 220 - 460 ms | 380 ms | 0.0055 | 1.1164 | 210 - 990 ms | 600 ms | 0.0001 | 1.1962 |  |
 500 - 680 ms | 570 ms | 0.0129 | 0.9103 |  | | | |

  

b) **Forward Concealed**  
trained on Incidental Recognition: incidental familiar vs. incidental unfamiliar  
tested on Concealed Knowledge: concealed familiar vs. true unfamiliar

|  | time window | peak latency | cluster *p* | peak Cohen's *d* |  | | | |  |
| **all electrodes** | 220 - 780 ms | 260 ms | 0.0001 | 0.983 |  | | | |  |
|  | | | | | | | | |  |
|  | **left hemisphere** | | | | **right hemisphere** | | | |  |
|  | time window | peak latency | cluster *p* | peak Cohen's *d* | time window | peak latency | cluster *p* | peak Cohen's *d* |  |
| **anterior** | 230 - 460 ms | 400 ms | 0.0006 | 1.5832 | 230 - 450 ms | 260 ms | 0.0033 | 0.8259 |  |
| **central** | 220 - 660 ms | 340 ms | 0.0002 | 1.3714 | 200 - 780 ms | 250 ms | 0.0004 | 1.07 |  |
  | | | | 810 - 900 ms | 840 ms | 0.0448 | 0.7148 ||  | **posterior** | 210 - 750 ms | 250 ms | 0.0001 | 0.9202 | 180 - 990 ms | 260 ms | 0.0001 | 1.0002 |
 830 - 990 ms | 860 ms | 0.0125 | 0.7657 |  | | | |

  

c) **Reverse Acknowledged**  
trained on Concealed Knowledge: acknowledged familiar vs. true unfamiliar  
tested on Incidental Recognition: incidental familiar vs. incidental unfamiliar

|  | time window | peak latency | cluster *p* | peak Cohen's *d* |  | | | |  |
| **all electrodes** | 210 - 810 ms | 440 ms | 0.0001 | 1.1333 |  | | | |  |
 830 - 910 ms | 840 ms | 0.0346 | 0.88 |  | | | ||  |  | | | | | | | | |
|  | **left hemisphere** | | | | **right hemisphere** | | | |  |
|  | time window | peak latency | cluster *p* | peak Cohen's *d* | time window | peak latency | cluster *p* | peak Cohen's *d* |  |
| **anterior** | 210 - 290 ms | 250 ms | 0.0339 | 0.8621 | 220 - 810 ms | 430 ms | 0.0001 | 1.5219 |  |
 350 - 740 ms | 530 ms | 0.0003 | 1.0903 |  | | | ||  | **central** | 210 - 990 ms | 430 ms | 0.0001 | 2.1946 | 200 - 990 ms | 440 ms | 0.0001 | 2.7169 |
| **posterior** | 200 - 440 ms | 400 ms | 0.0045 | 0.8792 | 330 - 900 ms | 560 ms | 0.0001 | 1.1736 |  |
 500 - 720 ms | 580 ms | 0.0037 | 0.9695 |  | | | | 830 - 990 ms | 950 ms | 0.0179 | 0.7284 |  | | | |

  

d) **Reverse Concealed**  
trained on Concealed Knowledge: concealed familiar vs. true unfamiliar  
tested on Incidental Recognition: incidental familiar vs. incidental unfamiliar

|  | time window | peak latency | cluster *p* | peak Cohen's *d* |  | | | |
| **all electrodes** | 230 - 990 ms | 430 ms | 0.0002 | 1.1036 |  | | | |
|  | | | | | | | | |
|  | **left hemisphere** | | | | **right hemisphere** | | | |
|  | time window | peak latency | cluster *p* | peak Cohen's *d* | time window | peak latency | cluster *p* | peak Cohen's *d* |
| **anterior** | 330 - 790 ms | 420 ms | 0.0001 | 1.2808 | 220 - 990 ms | 540 ms | 0.0001 | 2.0084 |
| **central** | 200 - 990 ms | 430 ms | 0.0001 | 2.3273 | 210 - 990 ms | 540 ms | 0.0001 | 1.7505 |
| **posterior** | 220 - 780 ms | 260 ms | 0.0009 | 1.37 | 220 - 990 ms | 550 ms | 0.0001 | 1.1281 |
 820 - 990 ms | 960 ms | 0.0143 | 0.9173 |  | | | |

  

e) **Forward Lie**  
trained on Incidental Recognition: incidental familiar vs. incidental unfamiliar  
tested on Concealed Knowledge: concealed familiar vs. acknowledged familiar

|  |  |  |  |  |  |  |  |  |
| --- | --- | --- | --- | --- | --- | --- | --- | --- |
|  | time window | peak latency | cluster *p* | peak Cohen's *d* |  | | | |
| **all electrodes** |  | | | |  | | | |
|  | **left hemisphere** | | | | **right hemisphere** | | | |
|  | time window | peak latency | cluster *p* | peak Cohen's *d* | time window | peak latency | cluster *p* | peak Cohen's *d* |
| **anterior** |  | | | |  | | | |
| **central** |  | | | |  | | | |
| **posterior** |  | | | | 240 - 340 ms | 260 ms | 0.0313 | -0.6895 |

  

f) **Reverse Lie**  
trained on Concealed Knowledge: concealed familiar vs. acknowledged familiar  
tested on Incidental Recognition: incidental familiar vs. incidental unfamiliar

|  | time window | peak latency | cluster *p* | peak Cohen's *d* |  | | | |  |
| **all electrodes** |  | | | |  | | | |  |
|  | | | | | | | | |  |
|  | **left hemisphere** | | | | **right hemisphere** | | | |  |
|  | time window | peak latency | cluster *p* | peak Cohen's *d* | time window | peak latency | cluster *p* | peak Cohen's *d* |  |
| **anterior** | 400 - 460 ms | 420 ms | 0.0195 | 1.0964 | 410 - 470 ms | 430 ms | 0.0273 | 0.8846 |  |
 490 - 590 ms | 520 ms | 0.0155 | 0.7872 |  | | | | 860 - 940 ms | 890 ms | 0.0151 | -1.045 |  | | | ||  | **central** | 380 - 710 ms | 520 ms | 0.0001 | 1.452 | 380 - 710 ms | 530 ms | 0.0003 | 1.5563 |
|  | **posterior** | 540 - 600 ms | 590 ms | 0.0068 | 1.2404 | 220 - 280 ms | 250 ms | 0.0348 | -1.0268 |
  | | | | 310 - 360 ms | 330 ms | 0.0253 | -1.3006 |

  
