## Supplementary Table 3 for "Characterizing the shared signals of face familiarity: long-term acquaintance, voluntary control, and concealed knowledge": SupplementaryTable_3.html

  

a) **incidental familiar**

|  | time window | peak latency | cluster *p* | peak Cohen's *d* |  | | | |  |
| **all electrodes** | 220 - 990 ms | 480 ms | 0.0001 | 2.1349 |  | | | |  |
|  | | | | | | | | |  |
|  | **left hemisphere** | | | | **right hemisphere** | | | |  |
|  | time window | peak latency | cluster *p* | peak Cohen's *d* | time window | peak latency | cluster *p* | peak Cohen's *d* |  |
| **anterior** | 220 - 310 ms | 250 ms | 0.0232 | 1.1582 | 210 - 980 ms | 460 ms | 0.0001 | 1.7174 |  |
 330 - 780 ms | 440 ms | 0.0003 | 1.3853 |  | | | | 800 - 950 ms | 930 ms | 0.0143 | 0.8183 |  | | | ||  | **central** | 220 - 990 ms | 430 ms | 0.0001 | 2.0151 | 210 - 990 ms | 440 ms | 0.0001 | 2.5563 |
|  | **posterior** | 190 - 990 ms | 510 ms | 0.0001 | 1.6651 | 220 - 990 ms | 490 ms | 0.0001 | 1.7859 |

  

b) **incidental unfamiliar**

|  | time window | peak latency | cluster *p* | peak Cohen's *d* |  | | | |  |
| **all electrodes** | 210 - 990 ms | 470 ms | 0.0001 | 2.1979 |  | | | |  |
|  | | | | | | | | |  |
|  | **left hemisphere** | | | | **right hemisphere** | | | |  |
|  | time window | peak latency | cluster *p* | peak Cohen's *d* | time window | peak latency | cluster *p* | peak Cohen's *d* |  |
| **anterior** | 240 - 790 ms | 460 ms | 0.0001 | 1.7911 | 240 - 980 ms | 550 ms | 0.0001 | 2.2654 |  |
 810 - 930 ms | 900 ms | 0.0238 | 0.5427 |  | | | ||  | **central** | 200 - 990 ms | 480 ms | 0.0001 | 2.0585 | 220 - 990 ms | 470 ms | 0.0001 | 2.3604 |
| **posterior** | 220 - 990 ms | 470 ms | 0.0001 | 1.9941 | 210 - 990 ms | 470 ms | 0.0001 | 1.9351 |  |

  

c) **concealed familiar**

|  | time window | peak latency | cluster *p* | peak Cohen's *d* |  | | | |
| **all electrodes** | 230 - 430 ms | 260 ms | 0.0004 | 0.8085 |  | | | |
|  | | | | | | | | |
|  | **left hemisphere** | | | | **right hemisphere** | | | |
|  | time window | peak latency | cluster *p* | peak Cohen's *d* | time window | peak latency | cluster *p* | peak Cohen's *d* |
| **anterior** |  | | | |  | | | |
| **central** | 240 - 370 ms | 340 ms | 0.0023 | 0.8554 | 200 - 410 ms | 250 ms | 0.0008 | 1.2055 |
| **posterior** | 220 - 430 ms | 320 ms | 0.0004 | 0.8124 | 190 - 420 ms | 260 ms | 0.0005 | 1.0363 |
 830 - 910 ms | 840 ms | 0.0141 | 0.6949 |  | | | |

  

d) **acknowledged familiar**

|  | time window | peak latency | cluster *p* | peak Cohen's *d* |  | | | |
| **all electrodes** | 390 - 670 ms | 550 ms | 0.0008 | 1.0063 |  | | | |
|  | | | | | | | | |
|  | **left hemisphere** | | | | **right hemisphere** | | | |
|  | time window | peak latency | cluster *p* | peak Cohen's *d* | time window | peak latency | cluster *p* | peak Cohen's *d* |
| **anterior** |  | | | |  | | | |
| **central** | 540 - 640 ms | 570 ms | 0.0422 | 0.5217 | 370 - 890 ms | 430 ms | 0.0004 | 0.8501 |
| **posterior** | 360 - 430 ms | 380 ms | 0.0488 | 0.7406 | 350 - 490 ms | 430 ms | 0.0137 | 0.6545 |
 540 - 620 ms | 570 ms | 0.0418 | 0.7226 | 530 - 700 ms | 560 ms | 0.0101 | 0.7658 |

  

e) **true unfamiliar**

|  | time window | peak latency | cluster *p* | peak Cohen's *d* |  | | | |  |
| **all electrodes** | 210 - 850 ms | 640 ms | 0.0001 | 1.2789 |  | | | |  |
 920 - 990 ms | 940 ms | 0.0439 | 0.9376 |  | | | ||  |  | | | | | | | | |
|  | **left hemisphere** | | | | **right hemisphere** | | | |  |
|  | time window | peak latency | cluster *p* | peak Cohen's *d* | time window | peak latency | cluster *p* | peak Cohen's *d* |  |
| **anterior** | 240 - 610 ms | 410 ms | 0.0005 | 1.0797 | 300 - 620 ms | 420 ms | 0.0013 | 1.4849 |  |
  | | | | 640 - 720 ms | 660 ms | 0.0481 | 0.7139 ||  | **central** | 210 - 650 ms | 570 ms | 0.0001 | 0.9624 | 220 - 990 ms | 600 ms | 0.0001 | 1.5303 |
| **posterior** | 210 - 460 ms | 390 ms | 0.0025 | 1.1602 | 180 - 990 ms | 640 ms | 0.0001 | 1.4489 |  |
 520 - 680 ms | 570 ms | 0.0135 | 0.8457 |  | | | |

  

f) **acknowledged familiar - true unfamiliar**

|  | time window | peak latency | cluster *p* | peak Cohen's *d* |  | | | |
| **all electrodes** | 230 - 290 ms | 250 ms | 0.0467 | -0.733 |  | | | |
|  | | | | | | | | |
|  | **left hemisphere** | | | | **right hemisphere** | | | |
|  | time window | peak latency | cluster *p* | peak Cohen's *d* | time window | peak latency | cluster *p* | peak Cohen's *d* |
| **anterior** |  | | | |  | | | |
| **central** | 280 - 360 ms | 340 ms | 0.0235 | -0.7144 | 240 - 360 ms | 330 ms | 0.0045 | -0.9416 |
| **posterior** | 280 - 440 ms | 280 ms | 0.0014 | -0.5573 | 160 - 360 ms | 300 ms | 0.0008 | -0.9558 |
  | | | | 600 - 660 ms | 640 ms | 0.0347 | -0.9277 |

  

g) **concealed familiar - true unfamiliar**

|  | time window | peak latency | cluster *p* | peak Cohen's *d* |  | | | |  |
| **all electrodes** | 430 - 660 ms | 440 ms | 0.0012 | -0.7997 |  | | | |  |
 730 - 790 ms | 770 ms | 0.0445 | -0.6286 |  | | | ||  |  | | | | | | | | |
|  | **left hemisphere** | | | | **right hemisphere** | | | |  |
|  | time window | peak latency | cluster *p* | peak Cohen's *d* | time window | peak latency | cluster *p* | peak Cohen's *d* |  |
| **anterior** | 410 - 590 ms | 420 ms | 0.0034 | -0.6199 | 400 - 490 ms | 430 ms | 0.0183 | -0.6642 |  |
  | | | | 510 - 600 ms | 570 ms | 0.0094 | -0.9825 ||  | **central** | 510 - 620 ms | 570 ms | 0.0112 | -0.7386 | 410 - 470 ms | 430 ms | 0.0439 | -0.7894 |
  | | | | 500 - 730 ms | 540 ms | 0.0016 | -1.0389 || **posterior** |  | 360 - 440 ms | 380 ms | 0.0372 | -0.642 | 490 - 730 ms | 590 ms | 0.0012 | -0.9111 |
 530 - 640 ms | 560 ms | 0.0137 | -0.7851 |  | | | |

  

h) **concealed familiar - acknowledged familiar**

|  |  |  |  |  |  |  |  |  |
| --- | --- | --- | --- | --- | --- | --- | --- | --- |
|  | time window | peak latency | cluster *p* | peak Cohen's *d* |  | | | |
| **all electrodes** |  | | | |  | | | |
|  | **left hemisphere** | | | | **right hemisphere** | | | |
|  | time window | peak latency | cluster *p* | peak Cohen's *d* | time window | peak latency | cluster *p* | peak Cohen's *d* |
| **anterior** |  | | | |  | | | |
| **central** |  | | | |  | | | |
| **posterior** |  | | | |  | | | |

  
